# *F*_*ST*_ and genetic diversity in an island model with background selection

**DOI:** 10.1101/2024.03.15.585154

**Authors:** Asad Hasan, Michael C. Whitlock

## Abstract

Background selection, by which selection on deleterious alleles reduces diversity at linked neutral sites, influences patterns of total neutral diversity, *π*_*T*_, and genetic differentiation, *F*_*ST*_, in structured populations. The theory of background selection may be split into two regimes: the *background selection regime*, where selection pressures are strong and mutation rates are sufficiently low such that deleterious alleles are at a deterministic mutation-selection balance, and the *interference selection regime*, where selection pressures are weak and mutation rates are sufficiently high that deleterious alleles accumulate and interfere with another, leading to selective interference. Previous work has quantified the effects of background selection on *π*_*T*_ and *F*_*ST*_ only for deleterious alleles in the *background selection regime*. Furthermore, there is evidence to suggest that migration reduces the effects of background selection on *F*_*ST*_, but this has not been fully explained. Here, we derive novel theory to predict the effects of migration on background selection experienced by a subpopulation, and extend previous theory from the *interference selection regime* to make predictions in an island model. Using simulations, we show that this theory best predicts *F*_*ST*_ and *π*_*T*_. Moreover, we demonstrate that background selection from weakly deleterious alleles may generate minimal increases in *F*_*ST*_, because migration reduces correlated effects on fitness over generations within subpopulations. However, we show that background selection may still cause substantial reductions in *π*_*T*_, particularly for metapopulations with a larger effective population size. Our work further extends the theory of background selection into structured populations, and suggests that background selection will minimally confound locus-to-locus *F*_*ST*_ scans.

**Author Summary:** Most mutations that affect fitness incur deleterious effects and are ultimately removed via natural selection. Consequently, nearby neutral variants may also experience the effects of selection; this is termed background selection. Background selection greatly influences patterns of genetic diversity both between and within populations among virtually all extant species, and is therefore of great interest to geneticists. Previous models of background selection have been primarily restricted to populations with completely random mating. However, it is well known that most natural populations exhibit some form of spatial structure. Here, we explore the effects of background selection in spatially structured populations, and we find that migration between subpopulations may attenuate the effects of background selection acting to increase genetic differentiation among populations. We derive novel theory to account for this effect by considering that individuals with deleterious alleles may migrate out of a local subpopulation prior to being removed by the population via selection. Our work demonstrates that, when migration rates are high, background selection does not substantially influence genetic differentiation among populations. Despite this, we find that background selection may greatly decrease genetic diversity within subpopulations and in the whole metapopulation.

## Introduction

Background selection is the process by which selection on deleterious alleles also affects diversity at linked sites [1]. Background selection reduces genetic diversity, as individuals carrying deleterious mutations are selected out of the population, and ultimately do not contribute to neutral diversity [1-3]. Due to the ubiquity of deleterious mutations [4], background selection occurs in all species and its effects have been estimated in several species including humans [5-8], fruit flies [9-10], and various other eukaryotes [11]. Given that background selection influences patterns of genetic variation and the site frequency spectrum [12-14], it may bias demographic inference or inferences of other selective forces [15]. Thus, there is a general recognition of the effects of background selection on patterns of diversity, and efforts are underway to incorporate background selection into models of genomic diversity [7, 10, 16].

The strength of background selection is quantified using the metric *B*, the ratio of the amount of genetic diversity at a neutral locus as a proportion of the expected diversity in the absence of background selection. In other words,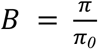, where *π* refers to observed neutral diversity and *π*_0_ refers to neutral diversity expected in the absence of selection on linked sites; π_0_≈4*N*_*e*_*µ* in a diploid population under an infinite-sites model [17] (where *N*_*e*_ is the effective population size and *µ* is the neutral per base-pair mutation rate). Generally, we can think of purifying selection as introducing positive correlations between generations in allele frequency change, amplifying the variance in reproductive success (and therefore genetic drift) experienced by linked neutral sites, further reducing their *N*_*e*_ beyond a simple mutation-drift model [18-19]. Individuals with deleterious mutations are less likely to have offspring, and so are their offspring as they may also carry deleterious mutations. The amplification of variance in reproductive success of neutral alleles due to their association with deleterious alleles is therefore captured by *B*.

Most natural populations exhibit some degree of population structure [20], and this influences patterns of genetic diversity. In structured populations, *F*_*ST*_ [21] is a widely used population genetic metric among plant/animal breeders, conservationists, and others [22] to identify sites subject to spatially varying selection (in the case of *F*_*ST*_ outlier tests), the demographic histories of subpopulations, and rates of local inbreeding. It can be defined as 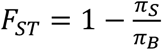, where *π*_S_ is the average proportion of pairwise differences between haplotypes within a subpopulation and *π*_B_ is the average proportion of pairwise differences between haplotypes from different subpopulations [23]. *F*_*ST*_ can also be defined in terms of the standardized variance in allele frequencies among subpopulations, or in terms of mean coalescence times for alleles within subpopulations versus the whole population [24].

Previous work has illustrated the effects of background selection on neutral diversity patterns in structured populations. [25-28]. Given that recombination rates are variable across the genome, background selection also varies across the genome. This generates locus-to-locus variation in *N*_*e*_ and *F*_*ST*_, which has been observed in human populations [6]. Locally beneficial alleles are expected to exhibit greater *F*_*ST*_ values relative to neutrality, as they would exhibit greater variance in frequency between subpopulations relative to neutral alleles. However, locus-to-locus variation in background selection may distort *F*_*ST*_ values for neutral loci as well, confounding scans for locally adapted alleles and potentially increasing the false-positive rate. Work by Matthey-Doret and Whitlock has shown this effect to be weak, particularly when mean *F*_*ST*_ values are low [28]. This work also suggested an effect of migration to increase *B* (reduce the effect of background selection on *N*_*e*_ and *π*). In other words, when migration rates are sufficiently high, it is possible some of the effects of background selection are not realized in a local population before the allele is lost to selection, which may weaken the confounding effect of background selection on *F*_*ST*_. In this paper, we investigate how varying migration rates influence the effect of background selection on *F*_*ST*_ and total diversity of a metapopulation, *π*_*T*_.

Theoretical studies suggest that the effects of background selection models can be generally divided into two regimes: the *background selection regime* and *interference selection regime* [29]. In the *background selection regime*, the key assumptions are that the effective strength of selection (i.e. the product of the effective population size multiplied by the heterozygous selection coefficient) |*N*_*e*_*t*| > 1 and mutation rate, *µ*, is small such that the deleterious alleles are introduced infrequently and selected out effectively. Here, the effects of background selection can be modeled following the assumptions of a deterministic mutation-selection balance, and dynamics at deleterious loci can be assumed to be independent from one another. Background selection was first modeled assuming an infinitely large diploid population where deleterious loci are at mutation-selection balance in a non-recombining genome. In this case the fraction of deleterious mutation-free gametes could be modeled using a Poisson distribution with mean 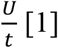 where *U* is the gametic genomic deleterious mutation rate and *t* is the strength of selection against deleterious mutations in heterozygotes. Soon after, this model was extended to consider recombination [2-3].

On the other hand, when selection against deleterious alleles is weak (i.e., |*N*_*e*_*t*| < 1), recombination rates are low or zero (*r* → 0), and the per base-pair mutation rate, *µ*, sufficiently high, deleterious alleles are introduced frequently and inefficiently removed by selection. Here, weakly deleterious alleles accumulate along the genome, and the effects of selection at a single site is now influenced by selection at other linked sites, leading to the “Hill-Robertson effect” or selective interference [30]. In this *interference selection regime*, assumptions of a deterministic mutation-selection balance fail to predict the quantitative effects of background selection, as deleterious alleles (|*N*_*e*_*t*| < 1) exhibit stochastic fluctuations in frequency due to the weak efficacy of selection and interactions between one another. In this regime, models cannot assume independence among selected sites and must consider multiple selected sites in tight linkage with one another [31]. Work by Good et al. [29] has shown that the effects of many weakly deleterious alleles in this interference selection regime can be approximated by fewer, strongly deleterious alleles in the background selection regime that generate equal variance in fitness. This model circumvents the issue of selective interference to accurately predict *B* in panmictic populations for both the *background selection* and *interference selection regimes*, but has not been applied to make predictions about patterns of genetic diversity in structured populations.

In models of structured populations under background selection, the equilibrium effects of background selection on *F*_*ST*_ may be approximated by multiplying the local population size of a subpopulation by *B* [27]. This is intuitive, as *B* represents the effects of background selection on the rate of genetic drift on neutral loci, with smaller values of *B* corresponding to greater amounts of genetic drift; thus, a greater effect of background selection on neutral loci will lead to a reduced local effective population size, *N*_*e, local*_. In a haploid two-island model, Zeng and Corcoran [27] predicted that

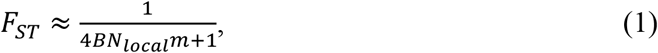

where *N*_*local*_ is the local population size, and *m* is the total migration rate. Note that *BN*_*local*_ = *N*_*e, local*_, and that this equation is consistent with the more general diploid finite island model prediction of 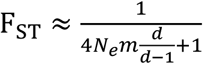, where *d* is the number of demes [32]. This approximation has been shown to work well to predict *F*_*ST*_ in the deterministic *background selection regime*, in cases with strong selection and low migration rates [27]. However, no current study provides a solution to the effects of background selection on *F*_*ST*_ for weakly deleterious alleles in the *interference selection regime*. This represents a large gap in the literature, as both the theory [33] and empirical estimates of the distribution of fitness effects (reviewed in Ref. [4]) suggest that most mutations incur weakly deleterious effects on fitness. Furthermore, we lack theory predicting the effects of migration on background selection, and the effects of background selection on total diversity, *π*_*T*_, in structured populations. Given that spatial structure is a common feature of natural populations, we require theory to better account for the role of migration, and its role in shaping *π*_*T*_.

The overall goal of this paper is therefore to better understand the effects of background selection in structured populations using simulations, and to further develop the theory to predict *π*_*T*_ and *F*_*ST*_ under background selection. Our simulations show that the effects of background selection from weakly deleterious alleles (in the *interference selection regime*) on *F*_*ST*_ are relatively weak, and that the effects of background selection on *F*_*ST*_ are attenuated by migration, approaching *F*_*ST*_ values expected under neutrality. We derive novel background selection theory to account for this migration effect. We also combine theory about background selection from the *interference selection regime* in panmictic populations [29] with what is known about how background selection affects *F*_*ST*_ [27] to best estimate *F*_*ST*_ and *π*_*T*_ in both the *interference selection* and *background selection regimes* under various strengths of selection, recombination rates, and migration rates. Lastly, we demonstrate that, while the effects of background selection on *F*_*ST*_ may be relatively weak, background selection can substantially reduce total *π*_*T*_ in a large metapopulation, and that we can predict this effect reasonably well. Our work extends the theory of background selection from panmictic populations to structured populations using an island model and lends further support for an insignificant confounding role of background selection in *F*_*ST*_ scans for populations where *F*_*ST*_ is relatively low on average. Our work also suggests an under-appreciated role for background selection from weakly deleterious alleles in reducing total diversity in sizable metapopulations.

## Theory

Zeng and Corcoran [27] predicted that the effects of *F*_*ST*_ in an island model would be predictable from modifying the effective size of a subpopulation, *N*_*local*_, by multiplying times *B*, the *N*_*e*_*/N* ratio created by background selection. We show here that migration lessens the effects of background selection on the amount of genetic differentiation caused by local drift relative to predictions of *B* based on earlier work (e.g. Ref [3]). Then, we derive the expected *F*_*ST*_ in an island model with background selection and a finite number of demes, when migration follows genetic drift.

## Derivation of *B* with ‘migration effect’

In this section, we use the approach from the Appendix of Ref. [34] to account for how migration affects *B* in structured populations.

In a metapopulation, suppose we have an island model with *d* subpopulations, each with *N*_*local*_ diploid individuals. Deleterious mutations appear at site *i* (and eventually achieve mutation-selection balance) with per-base pair mutation rate *µ*_*i*_ and heterozygous selection coefficient *t*_*i*_ (fitness is multiplicative across sites). Recombination between neutral and deleterious sites occurs at rate *r*_*i*_. Recombination changes genotype frequencies only if individuals do not die from selection following the recombination event; hence, the effective recombination rate is 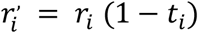. Migration occurs at rate *m*, with *m* fraction of individuals in each deme migrating out each generation and being replaced by immigrant individuals which come from a randomly chosen other subpopulation.

In order to predict the effect of background selection on *F*_*ST*_, we need to understand the effect of background selection on the effective population size of a single deme: the *local* effective population size, *N*_*e, local*_. The local effective population size will be affected by the variance in reproductive success of neutral alleles created by their association with deleterious mutations, just as in any undivided population. However, if an individual with the deleterious mutation migrates to another deme, any subsequent effects on the reproductive success of linked neutral sites will not affect *N*_*e, local*_. Therefore, we are interested in the variance in reproductive success of the neutral allele due to its association with the deleterious allele that affects the local distribution of reproductive success. Such variance in reproductive success will influence the local effective population size, *N*_*e, local*_, and ultimately neutral diversity, *π*_*local*_ and *B*_*local*_, within the subpopulation.

There are four factors that will influence the the effect of background selection on *N*_*e, local*_: the deleterious mutation rate per base-pair, *µ*_*i*_, the heterozygous selection coefficient, *t*_*i*_, of deleterious mutations, the recombination rate between the deleterious and neutral allele, *r*_*i*_, and the migration rate, *m*. The deleterious mutation rate, *µ*_*i*_, will affect the frequency of introduction of deleterious alleles, thus directly influencing the variance in reproductive success of neutral alleles and ultimately *B*_*local*_. The selection coefficient of a deleterious allele will influence the number of generations it remains in the population, with higher selection coefficients leading to more rapid purging and lower selection coefficients leading to patterns of interference among deleterious alleles. The recombination rate between a deleterious and a neutral allele will influence the number of generations they remain in association. All of these previous factors are present in classical derivations of the effects of background selection in single populations. In addition, when tracking the local effects of background selection, we add the expectation that the migration rate will influence the association of a neutral allele and deleterious allele *within the focal subpopulation*, as the emigration of a deleterious haplotype from the subpopulation will end the local effect of the deleterious allele on the reproductive success of the neutral allele.

Following Ref. [34], we consider two cases corresponding to whether a neutral allele is in coupling (case 1) or repulsion (case 2) with a deleterious allele in a single diploid individual. In case (1), the association between the neutral and deleterious allele can be ended by recombination, selection, or migration to another subpopulation, so the fraction of descendants of the neutral allele in the same subpopulation that carry the deleterious allele *i, n* generations later, is 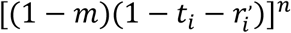. Taking the sum from *n =* 0 to ∞,

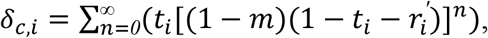

the expected net reduction in frequency of the neutral allele in this subpopulation over all generations is

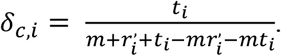

In case (2), the deleterious allele has an average reduction in frequency of *t*_*i*_ in the initial generation because of the presence in the same individual of the deleterious allele in repulsion, which only affects the local drift if that individual has not migrated. During this generation, there is a *r*^’^ probability that the deleterious allele recombines onto the background of the neutral variant. If it does so, then the future reduction in fitness while it is in the same local population is the same as in case 1. Therefore

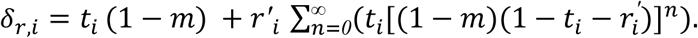

Taking the sum from *n = 0* to ∞, we find that the expected net reduction in neutral allele frequency here is

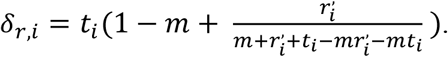

The total additive genetic variance at the neutral site over time, due to its associations with the deleterious allele through Case (1) and (2), increases the variance in reproductive success by Δ*V*_*i*_ ≈ *q*(*δ*_*c*_ + *δ*_*r*_) at the neutral site [34] beyond that of a mutation-drift model. Here, *q* is the frequency of the deleterious allele at mutation-selection balance. Given that 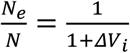, we may now estimate *N*_*e*_ at the neutral site experiencing background selection. Extending this to multiple deleterious sites in association with the neutral site at the local population,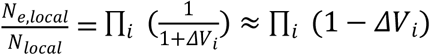 for small Δ*V* . Using Δ*V*_*i*_ ≈ *q*(*δ*_*a*_ + *δ*_*r*_ ), we find that

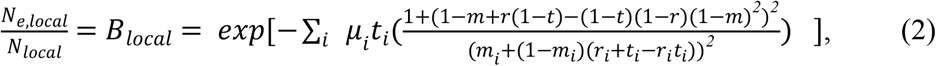

which for small values of *r*_*i*_ and *t*_*i*_ is approximately

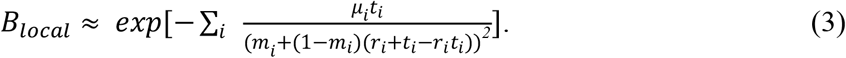

For a chromosome with total deleterious mutation rate *U* and total recombination rate *R*, the methodology used by Hudson and Kaplan [2] can be used to find that the above formula is well approximated by 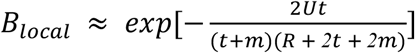. This requires a few assumptions: the neutral site is embedded in the center of a large region subject to deleterious mutations, *t* and *m* are fixed, and *r*_*i*_ is sufficiently small such that it is additive over sites. When *m* = 0, this formula reduces to 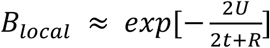.

For our predictions in the simulations, however, note that we do not directly employ Eq. 3 to predict *B*. Although it serves as a useful approximation, it ignores higher-order corrections (similar to classic theory from the *background selection regime*). Furthermore, for parameters such that we are in the *interference selection regime*, we utilize the methodology of Good et al. [29] in order to output parameters suitable for our derivation, thus allowing us to make accurate predictions when interference is common. For more details, see *Calculation of B* in the *Methods*.

## Prediction of *FST*

Previous work on background selection in structured populations has demonstrated that the effects of background selection in an island model can be approximated as a reduction in local effective population sizes proportional to the strength of background selection [25, 27]. In order to estimate and predict *F*_*ST*_, we use the definition of Hudson et al. [35], where

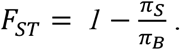

We show in the Appendix (Eq. A4) that, using the neutral expectation of *F*_*ST*_ in an island model, predicted *F*_*ST*_ values under background selection were derived as follows:

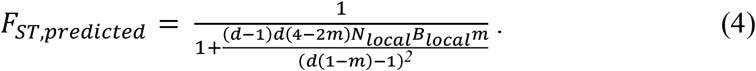

Here, *m* is the total proportion of immigrants into a deme per generation, *d* is the number of demes, and *B* is a measure of background selection experienced by the subpopulation. Our equation differs from that of Ref. [32] because, in SLiM (the evolutionary simulation framework used; [36]), the order of events is such that offspring production occurs first (genetic drift) followed by migration and then *F*_*ST*_ estimation [37], and because we consider terms that are *m*^*2*^ or higher.

We note that there are multiple definitions of *F*_*ST*_ that have been previously used. For instance, the equation used to estimate *F*_*ST*_ in our simulations uses the ratio of *π*_*S*_ and *π*_*B*_, but an alternative definition uses the ratio of *π*_*S*_ and *π*_*T*_, the total population diversity. This is equivalent to *G*_*ST*_, and is defined as:

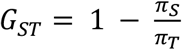

[38]. Using the neutral expectation, *G*_*ST*_ in a finite island model experiencing background selection with migration following drift and then *F*_*ST*_ estimation is derived in the Appendix (Eq. A5) to be:

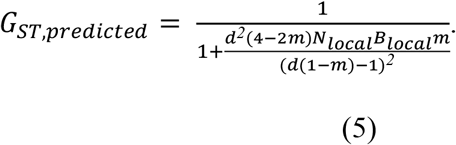

## Results

We use a simulation-based approach to investigate the effects of background selection in the *interference selection* and *background selection regimes* under varying migration rates. In SLiM v3.7.1 [36], we run simulations of a metapopulation using an island model consisting of *N*_*global*_ diploid individuals split equally into *d* demes. Population size is held constant, and generations are non-overlapping. Each generation, a total fraction of *m* individuals in each deme are replaced by immigrants, equally partitioned from each other deme. Genomes are composed of *L*_*neutral*_ neutral sites, surrounded on each side by selected sites for a total of *L*_*selected*_ subject to deleterious mutation. Recombination occurs between the neutral region and selected sites and between selected sites at total rate *R*. Deleterious mutations are introduced to gametes at fixed rate *U*, with fixed selection coefficients against heterozygotes *t*. Fitness is multiplicative across sites. More details are given in the *Methods* section.

### Background selection in structured populations and prediction by previous models

For our low migration cases (Fig. 1A & C), *F*_*ST*_ under neutrality was 0.50, thus representing populations with very high genetic differentiation. For our high migration cases (Fig. 1B & D), *F*_*ST*_ under neutrality was 0.004, representing populations with low genetic differentiation. Hereafter, we consider deleterious alleles with *N*_*local*_*t* ≤ 1 to be weakly deleterious, 1 < *N*_*local*_*t* < 7.5 to be moderately deleterious, and *N*_*local*_*t* ≥ 7.5 to be strongly deleterious. Given formulae from the theory of background selection following Good et al. [29], we expected the greatest increase in *F*_*ST*_ due to background selection at moderate strengths of selection. This is intuitive, as we expect for strongly deleterious mutations to be rapidly purged from the population, thus generating weak total variance in fitness due to their short duration in the population. Weakly deleterious mutations, on the other hand, may persist for many more generations in the population, but simply generate little total variance in fitness due to their individually weak selective effects. Moderately deleterious alleles, however, generate moderate variance in fitness and are not rapidly purged from the population; thus, we expected an intermediate maximum of *F*_*ST*_ with respect to strength of selection against deleterious alleles.

**Figure 1.**
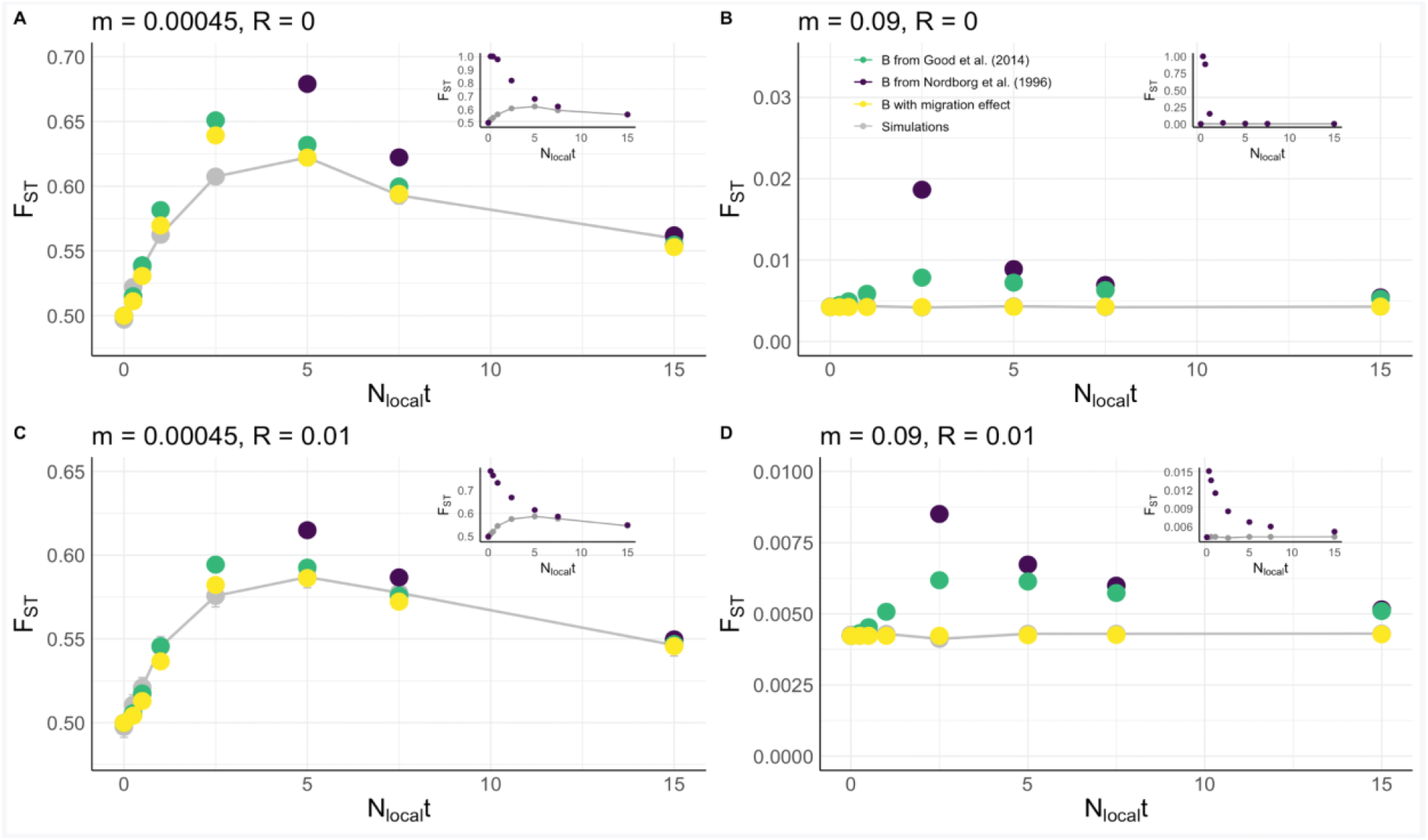
Accuracy of various methods in predicting *F*_*ST*_ with respect to the effective heterozygous strength of selection, *N*_*local*_*t*. (A) Forward-time simulations run with *m* = 0.00045 and *R* = 0, (B) *m* = 0.09 and *R* = 0, (C) *m* = 0.00045 and *R* = 0.01, and (D) *m* = 0.09 and *R* = 0.01. The gray dots connected by a line represent *F*_*ST*_ values from forward-time simulations, with 95% CIs. (These were quite small, and not always visible.) The yellow dots represent numerical predictions from our application of the Good et al. [29] method to structured populations with consideration of the ‘migration effect’ (see *Methods;* Eq. 7 & 8), and the green dots without its consideration (*Methods;* Eq. 7). The insets show the performance of classic background selection theory (purple dots; *Methods;* Eq. 6) to structured populations following Ref. [27]. All parameter combinations were simulated 750 times with local population size *N*_*local*_ = 500, fixed effective selection coefficients *N*_*local*_*t* = {0, 0.25, 0.5, 1, 2.5, 5, 7.5, 15}, *U* = 7x10^-3^, *L*_*selected*_ = 700, and for 250,000 generations. In many cases (especially in part D), the plot points for the simulations (gray dots) are hidden behind the points shown for theoretical expectations.

Indeed, we observed an inverted U-shape curve of *F*_*ST*_ under background selection, but only for our low migration cases (Fig. 1A & C); we later discuss this further. Considering only our low migration case, we observed that the effects of background selection on *F*_*ST*_ from weakly deleterious alleles are relatively weak (Fig. 1, gray dots). As expected, we also observed that the effect of background selection on *F*_*ST*_ is attenuated by recombination (Fig. 1C, *R* = 0.01, compared to A, *R* = 0); recombination introduces the opportunity for a neutral allele to recombine off of the background of a deleterious allele, thus rescuing it from eventual loss via linked selection.

Classic theory on the *background selection regime* (Ref. [2-3]; Fig. 1, purple dots) substantially overestimates the effects of background selection (underestimates *B*) for weakly deleterious alleles (see Ref. [29]) and therefore overestimates the effects on *F*_*ST*_ [27], with the discrepancy increasing for weaker strengths of selection (Fig. 1 insets, *cf*. purple to gray dots). As *t* approaches 0, the classical formula for *B* inaccurately predicts deleterious allele frequencies, and indeed those papers deriving the predictions are clear that the derivation applies to stronger effect mutations. The discrepancy between observation and prediction, however, is somewhat attenuated in the *R* = 0.01 cases (Fig. 1C & D, inset), presumably due to recombination reducing the magnitude of selective interference, a major driver of the discrepancy. Nevertheless, classic theory poorly predicts the effects of background selection on *F*_*ST*_ for weakly deleterious alleles, as expected.

On the other hand, the Good et al. [29] theory (Fig. 1, green dots) estimates *F*_*ST*_ well when alleles are weakly deleterious. However, there are two caveats. The theory does not accurately predict *F*_*ST*_ when *N*_*local*_*t* ≈ 2.5, and in our high migration parameter sets (Fig. 1B & D). *N*_*local*_*t* = 2.5 represents our chosen boundary between the *interference selection* and *background selection regimes*, and the formalization of this boundary remains an unsolved problem. Thus, we expect some inaccuracy from the theory here (see *Calculation of B* in *Methods*). In the case of high migration rates, the Good et al. [29] theory predicts an inverted-U shaped curve, but we observe no effect of background selection on *F*_*ST*_, for all values of *t*. This lack of effect of background selection on *F*_*ST*_ under higher rates of migration has been observed previously [28]. In our derivation above, we hypothesized that migration acts to reduce the effect of background selection by reducing the total variance in fitness over time experienced by neutral sites under background selection. We discuss its predictions below.

### ‘Migration effect’ theory accurately predicts the effects of background selection on *F*_*ST*_

The application of our derivation of *B* with the ‘migration effect’ (Fig. 1, yellow dots) accurately predicts *F*_*ST*_ under various fixed strengths of selection, recombination rates (Fig. 1), and migration rates (Fig. 2). Moreover, usage of the results of Good et al. [29] in conjunction with the new theory (Eq. 7 & 8) predicts *F*_*ST*_ well under lower strengths of selection, in the *interference selection regime*. As expected, the ‘migration effect’ is particularly prominent at higher migration rates. Under our higher migration rate parameter set (Fig. 1B & 1D), we observe no effect of background selection on *F*_*ST*_, despite marked reductions in *π*_*T*_ (Fig. 3B & D). However, the Good et al. [29] method predicts an inverted–U shaped increase in *F*_*ST*_. This discrepancy in Fig. 1B between the Good et al. [29] method and observation for all strengths of selection is likely due to large differences in the magnitude of *m* and *t* in this parameter set, with *m >> t* for all *t*. Here, migration exports deleterious alleles from their original patch and prevents the correlated loss of fitness over generations from affecting the effective population size of a single deme.

**Figure 2.**
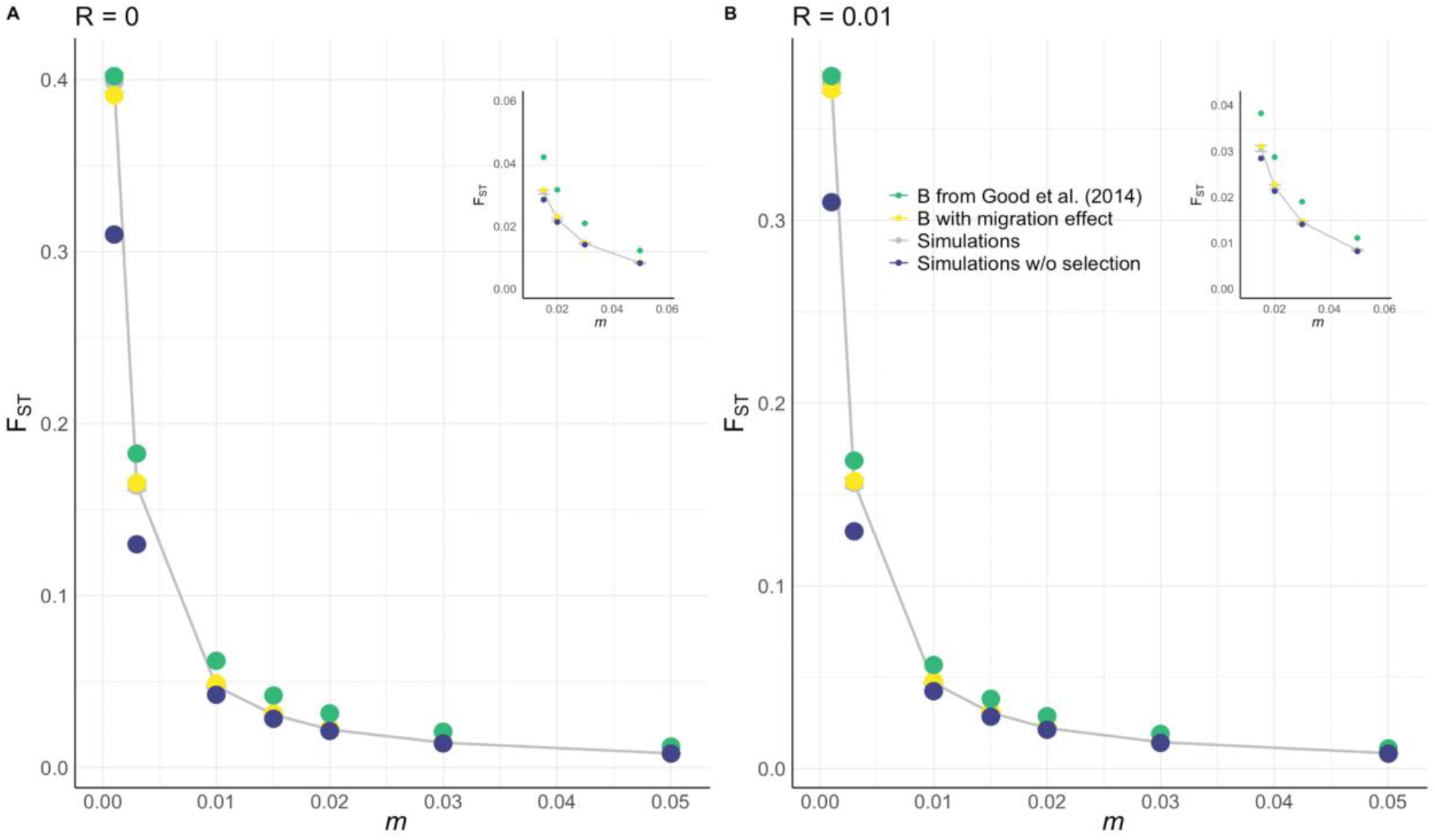
Inclusion and exclusion of migration effect in estimation of *F*_*ST*_ under background selection. Forward-time simulations, with deleterious alleles of selection coefficient *N*_*local*_*t* = 7.5, under varying rates of migration for (A) *R* = 0 and (B) *R* = 0.01. The gray dots represent observed *F*_*ST*_ and are connected by a line. The yellow dots represent our application of Good et al. [29] with the ‘migration effect’ (see *Methods;* Eq. 7 & 8), the green dots the Good et al. [29] method without the ‘migration effect’ (see *Methods*; Eq. 7), and the dark blue dots are *F*_*ST*_ from simulations run without deleterious alleles (*F*_*ST*_ under mutation-drift balance). The insets zoom in on cases for which *m* ≥ 0.015. All simulations were run 750 times with *N*_*local*_ = 500, *U* = 7x10^-3^, *L*_*selected*_ = 700, and for 250,000 generations.

**Figure 3.**
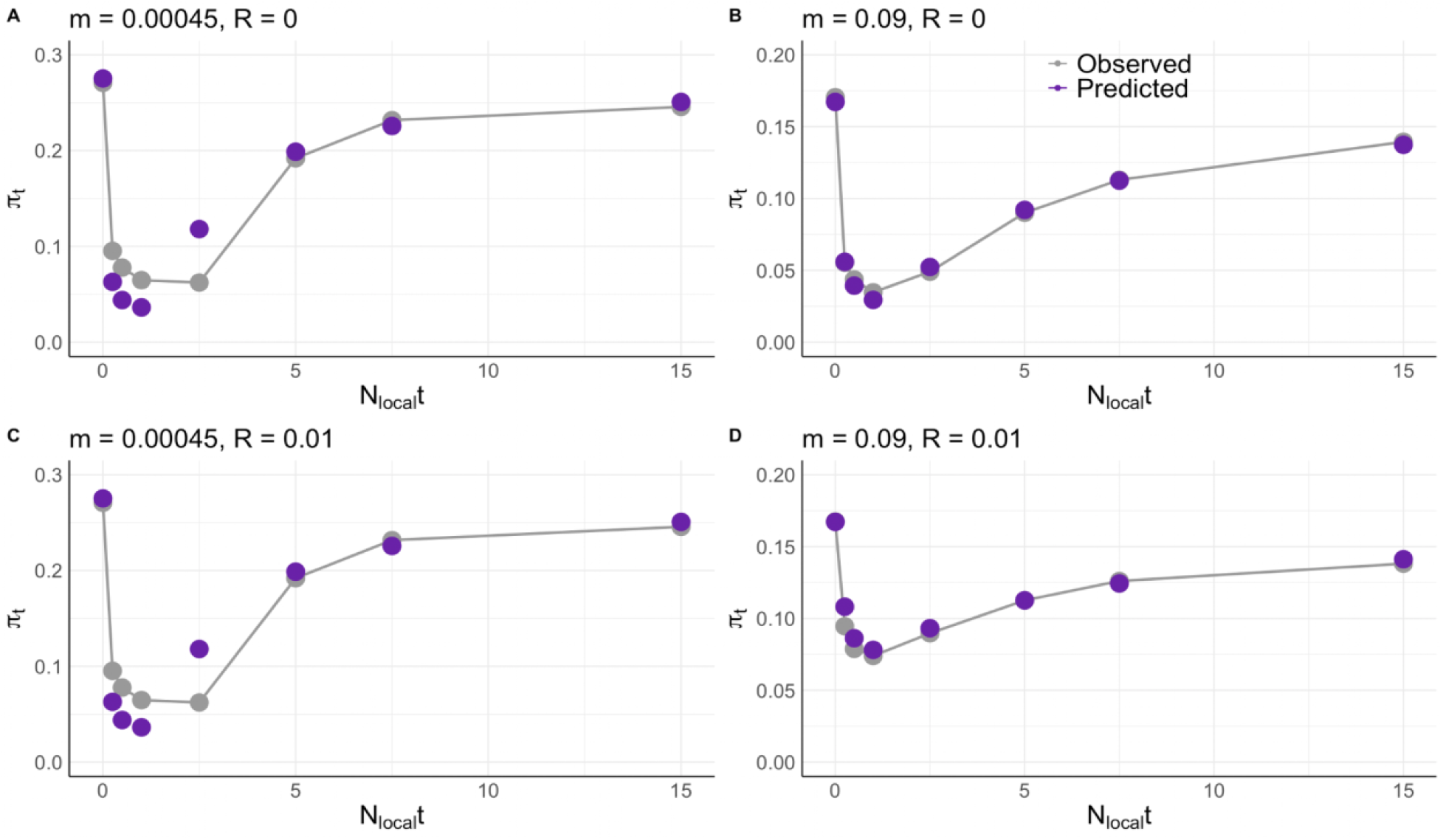
Predicted *π*_*T*_ versus observed *π*_*T*_ in a metapopulation. (A) Forward-time simulations run with *m* = 0.00045 and *R* = 0, (B) *m* = 0.09 and *R* = 0, (C) *m* = 0.00045 and *R* = 0.01, and (D) *m* = 0.09 and *R* = 0.01. The gray dots represent *π*_*T*_ from forward-time simulations using a 10-deme island model with *N*_*local*_ = 500, *N*_*local*_*t* = {0, 0.25, 0.5, 1, 2.5, 5, 7.5, 15}, *m* = {5x10^-5^, 1x10^-2^}, *r* = 0.005, and are connected by a line. The purple dots represent theoretical predictions of total diversity (see *Methods*) with incorporation of the migration effect to estimate *G*_*ST*,_ (*Theory* Eq. 5), *N*_*e, global*_, and therefore *π*_*T*_.

We explored the attenuating effect of varying rates of migration on background selection across multiple migration rates in Figure 2. For migration rates of *m =* 0.01 and higher, we find that *F*_*ST*_ values (Fig. 2, gray dots) strongly resemble *F*_*ST*_ values (Fig. 2, dark blue dots) expected under no background selection. Thus, our results suggest that the alignment of observed *F*_*ST*_ value to neutral prediction at low *F*_*ST*_ values may not be sufficient to predict the absence of background selection in empirical systems, as it may otherwise be explained by the presence of both background selection and weak population structure. Regardless, we expect background selection to weakly confound locus-to-locus *F*_*ST*_ scans when gene flow is sufficiently common.

A caveat to our predictions, however, is the lack of accurate prediction at the boundary between the *interference selection regime* and *background selection regime* [29]. This boundary was chosen somewhat arbitrarily prior to simulation, and we expect some deviation between prediction and observation as we approach this zone. However, this remains an unsolved gap in the background selection literature. This is seen for our simulations at *m =* 0.00045 and *R =* 0 (Fig. 1A), where we observe a substantial discrepancy between our best predictions and observation at *N*_*local*_*t =* 2.5, despite the model performing well for all other strengths of selection.

### Background selection may substantially reduce *π*_*T*_, and this is well predicted by theory

We found that background selection can cause substantial reductions in *π*_*T*_, despite weak effects on *F*_*ST*_. This effect is most pronounced from weakly deleterious alleles, for both low and high migration parameter sets, and is insignificantly influenced by the introduction of low rates of recombination. Previous work has suggested that selective interference is more common in larger populations (reviewed by Ref. [39]); given that in our simulations *N*_*e, global*_ is on the order of 10^4^, this is likely the case. We investigated this further and found that the Good et al. [29] method indeed predicts a greater reduction in *B* at lower strengths of selection for increasing population sizes (Supp. Fig. S1). Overall, the results from our parameter sets suggest that background selection against weakly deleterious alleles may cause up to 3-fold reductions in *π*_*T*_ compared to neutrality. Moreover, we demonstrate that we are able to predict the effects of background selection on *π*_*T*_ with reasonable accuracy. A caveat to our results was that *π*_*T*_ was poorly predicted for *N*_*e*_*t* ≤ 2.5 under our low migration parameter set (Fig. 2A & C), under-predicting *π*_*T*_ for *N*_*e*_*t* ≤ 1 and over-predicting *π*_*T*_ for *N*_*e*_*t =* 2.5. We note that a similar bias was observed in predictions of *π*_*S*_, thus not substantially influencing *F*_*ST*_ predictions (Fig. 1A & C). Since we observed this with our low migration parameter set and not our high migration parameter set, we believe this effect is not influenced by increasing migration. Instead, it likely reflects inaccuracy with the Good et al. [29] method, particularly since *π*_*T*_ is worst predicted at our chosen boundary of the *interference selection* and *background selection regimes*.

Under our low migration rate parameter set, we found that *π*_*S*_ was reduced by a fractionally larger margin than *π*_*T*_, resulting in the elevation of *F*_*ST*_ (Supp. Fig. S2). However, in our high migration rate parameter set, *π*_*S*_ and *π*_*T*_ were similarly reduced for all simulated values of *N*_*local*_*s*. For instance, we observed that, at *N*_*local*_*t* = 5 with *m* = 0.00045 and *R =* 0, *π*_*T*_ was reduced by a fraction of ∼0.30 and *π*_*S*_ by a fraction of ∼0.45. At *N*_*local*_*t* = 5 with *m =* 0.09 and *R =* 0, on the other hand, *π*_*T*_ and *π*_*S*_ were both reduced by a fraction of ∼0.47.

## Discussion

Previous theory of background selection in the *interference selection regime* has been thus far restricted to panmictic populations. This represents a substantial gap in the background selection literature, as deleterious alleles, a ubiquitous feature of extant organisms, are typically expected to be weakly deleterious, and therefore suited to the *interference selection regime*. Moreover, most natural populations exist within a metapopulation framework. Here, we have extended the theory of background selection from the *interference selection regime* in a panmictic population [29] to that of an island model, and have derived novel theoretical predictions of how migration influences the effects of background selection in structured populations. We tested our novel derivations using simulations, and found that it best predicts *F*_*ST*_ and *π*_*T*_ in an island model for all tested parameters.

In background selection theory, the association of a neutral allele to a deleterious allele increases its variance in fitness beyond that of a simple mutation-drift model, because the deleterious allele introduces additional between-generation correlations in fitness [18-19] of the neutral allele. In our derivation of background selection accounting for the ‘migration effect’, we propose that increased migration rates lead to weaker effects of background selection by reducing the total excess variance in fitness experienced by a neutral allele within a single subpopulation. The magnitude of this excess variance in fitness is contingent on several factors, including the number of generations the neutral allele remains in association with the deleterious allele [34]. Deleterious alleles with greater heterozygous selection coefficients (*t*) are purged more rapidly, thus reducing the number of generations they persist in a population. This effect explains why deleterious alleles with relatively strong selection coefficients cause relatively weak increases in *F*_*ST*_ (Fig. 1) and reductions in *π*_*T*_ (Fig. 3); they simply do not last enough generations to substantially influence reproductive success of an associated neutral allele. Moreover, the recombination of a neutral allele off of the background of a deleterious allele reduces the number of generations the deleterious allele remains in association with a neutral allele, also reducing the total excess fitness variance experienced by the neutral allele. The effects of variable selection coefficients and recombination rates have previously been incorporated into models of background selection. Here, we add the additional possibility that if a deleterious haplotype migrates out of a population, it no longer affects the change in local allele frequency, truncating the cross-generational correlations in reproductive success.

Our model provides accurate predictions of *F*_*ST*_ and *π*_*T*_ in an island model. Charlesworth previously showed, in simulations of a two-deme island model with *m* = 0.01, that background selection increases *F*_*ST*_ primarily by decreasing *π*_*S*_, with lower reductions in *π*_*T*_. This is intuitive under a lower migration rate, as we expect deleterious alleles in a subpopulation to be selected out prior to immigration. Thus, the effects of background selection are primarily experienced locally. Generally, we expect that when *m* << *t*, the effects of background selection on *F*_*ST*_ are similar to those predicted by single-population estimates of *B*. When *m >> t*, on the other hand, deleterious haplotypes may migrate out of the subpopulation prior to selection; thus, we expect a relatively weaker reduction in *π*_*S*_ and less of an increase in *F*_*ST*_.

Additionally, our results may also hold implications for the role of background selection in the surprisingly weak correlation between census population sizes and diversity, also known as Lewontin’s Paradox (reviewed in Ref. [40]). Previous theory has suggested a direct correlation between *N*_*e*_ and diversity, *π*, [17] that has not been observed in naturally occurring populations. There have been various proposed explanations for this, including the effects of linked selection and demographic history. Our simulations indicate that background selection from weakly deleterious alleles may lead to substantial drops in *π*_*T*_ in large metapopulations (*N*_*e*_ on the order of 10^4^ in our simulations), greater than that of deleterious alleles with moderate and strong selection coefficients. We expect this effect to become increasingly substantial as *N*_*e*_ increases, due to a greater accumulated density of deleterious alleles and selective interference [39]. Indeed, the method followed by Good et al. [29] predicts a greater decrease in *B* for weakly deleterious alleles as *N* increases (Fig. S1). Thus, background selection from weakly deleterious alleles may play an under-appreciated role in decreasing *π* in large, naturally occurring metapopulations.

Our study only looked at an island model. However, the effects of background selection on *F*_*ST*_ and *π*_*T*_ remain unexplored for other population subdivision models, such as ‘isolation-by-distance’ models with varying dimensionality. Additionally, the modulating effect of migration on the effect of background selection and *F*_*ST*_ remains unexplored in these other population subdivision models. Here, we also use various other simplifying assumptions, including fixed selection coefficients and a simple genomic architecture. We did not explore the effects of varying deleterious selection coefficients on *B*, but it is possible that the combination of both weakly and moderately deleterious alleles may result in more complex dynamics of background selection; this is of particular interest, given that we expect a much greater frequency of weakly deleterious alleles than strongly deleterious alleles [4]. The incorporation of variable selection coefficients, recombination rates, and placement of neutral sites into the theory remains an outstanding gap to be bridged.

In summary, we find that the effects of background selection from weakly deleterious alleles on *F*_*ST*_ are relatively weak, and attenuated by migration. We derive predictions of *B* accounting for the effect of variable migration rates, and demonstrate that these predictions accurately predict *F*_*ST*_ and *π*_*T*_ in an island model. Lastly, we show that, despite relatively small effects of background selection on *F*_*ST*_, background selection even from weakly deleterious alleles may result in considerable reductions in *π*_*T*_.

## Methods

### Simulations

We ran simulations in SLiM v3.7.1 [36] using a finite-island model consisting of 10 demes with a local diploid population size, *N*_*local*_ = 500 individuals, for a total of *N*_*global*_ = 5,000 individuals. To delineate the effects of background selection on *F*_*ST*_ and *π*_*t*_ under varying strengths of selection, we ran simulations under the following parameter set: *N*_*local*_*t* = {0, 0.25, 0.5, 1, 2.5, 5, 7.5, 15}, *U* = 7x10^-3^, *R* = {0, 0.01}, *r* = {0, 1.414x10^-5^}, and *m* = {0.00045, 0.09}, where *U* is the total gametic deleterious mutation rate, *r* is the per base-pair recombination rate, *R* is the total recombination rate (in Morgans), *t* is the selection coefficient against heterozygotes with a deleterious allele, and *m* is the total migration rate (the fraction of individuals in each deme that are replaced by immigrants from all other demes). To delineate the effects of varying migration rates on the effect of background selection on *F*_*ST*_, we also ran simulations under the following parameter set: *N*_*local*_*t* = 7.5, *U* = 7x10^-3^, *R* = {0, 0.01}, and *m =* {0.001, 0.003, 0.01, 0.015, 0.02, 0.03, 0.05}.

Fitness effects were multiplicative, such that *w*_*i*_ = (1 -*t*)^k^, where *w*_*i*_ is the survival rate, *t* is the selection coefficient against heterozygotes, and *k* is the number of deleterious alleles carried by the individual. The genome consisted of *L*_*neutral*_ = 10^4^ neutral sites embedded in a region surrounded on each side by 350 sites subject to deleterious mutation, adding up to a total of *L*_*selected*_ = 700 selected sites. This is equivalent to a genome consisting of a number of sites under selection upon which recombination occurred at total rate *R*, with a non-recombining neutral region embedded in the center. Each simulation was run 750 times to derive 95% CIs and for 250,000 (50*N*_*global*_) generations to achieve equilibrium. Data from the simulations were saved as tree sequence files, and neutral mutations were overlaid using *msprime*. Tree sequence data was analyzed using the *pyslim* and *tskit* packages in Python 3.9.12 to output *F*_*ST*_, *π*_*S*_, and *π*_*T*_.

We note that the parameter sets used for this study, although biologically realistic on average, are held fixed (e.g. selection coefficients, recombination rates), as our primary objectives were to study the limits of the theory of background selection instead of simulating realistic background selection in a biological system subject to variable parameters. Thus, we opted for parameter sets that would allow us to identify substantial and differentiable effects of background selection with minimal computing power under various strengths of selection, migration, and recombination.

### Calculation of *F*_*ST*_

*F*_*ST*_ in the simulations was estimated as:

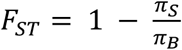

[23, 41], where *π*_*S*_ is the average proportion of pairwise differences between haplotypes sampled within a deme at a neutral locus and *π*_*B*_ is the average proportion of pairwise differences between haplotypes sampled between demes at a neutral locus. We filtered neutral alleles for which the minor allele frequency was less than 0.05 prior to calculating *F*_*ST*_.

### Calculation of *B*

For the case where deleterious mutations behave deterministically (i.e. the *background selection regime*), Nordborg et al. [3] predict *B*:

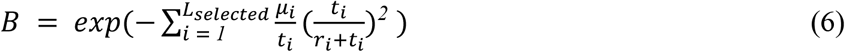

To derive our theoretical estimates of background selection in the *interference selection regime* in structured populations, we calculated *B* using the Python scripts supplied by Good et al. [29]. Briefly, Good et al. [29] found that the effects of background selection on neutral genetic diversity over an effectively non-recombining region, when interference is common, is determined by variance in fitness. They developed a method that approximates the effects of many weakly deleterious mutations in the *interference selection regime* as if there are fewer, strongly deleterious mutations in the *background selection regime* that generate equal variance in fitness. *B* is then estimated using these “effective parameters” in the *background selection regime* following classic background selection theory (i.e. Ref. [2-3]), although the formula used by Good et al. [29] slightly differs from that of previous formulae as they include higher order corrections. The formula used is:

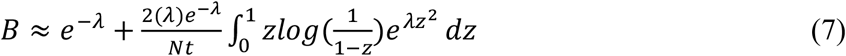

(Ref. [29] Eq. 2). Here, *t* is the selection coefficient against a deleterious allele in haploids (equivalent to the heterozygous selection coefficient in the case of diploids), *N* is the number of chromosome copies (i.e., population size for haploids and twice that for diploids), and 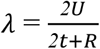 (described by Good et al. [29] as λ_*eff*_ ), where *R* is the total recombination rate in a selected region in which a neutral site is embedded in the center and *U* is the genome-wide gametic deleterious mutation rate. The parameters *U, t*, and *R* represent the “effective parameters” following use of the Good et al. [29] method. To predict *B* using our derivation including the effect of migration, we opt to use Eq. 2 of Good et al. [29], where instead

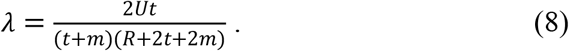

Therefore, we account for higher order corrections, migration, and selective interference by employing Eq. 7 & 8 to best predict *B* under varying *U, t, R*, and *m*.

We used the following cut-off for determining whether a particular parameter combination corresponded to the *interference selection regime*: 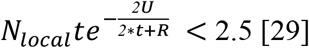. The precise formalization of the boundary between the *interference selection* and *background selection regimes* remains an unsolved problem in the background selection literature, but we found the aforementioned cut-off to be reasonable.

### Calculation and prediction of *π*_*T*_

In order to calculate *π*_*T*_ in a metapopulation, we must consider its global effective population size, *N*_*e, global*_. *N*_*e, global*_ is directly influenced by the degree of population structure, and, in the case of an island model, may be estimated as 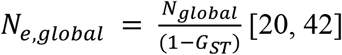. *G*_*ST*_ is the appropriate measure of population structure here, because it is standardized by total diversity due to its definitional use of *π*_*T*_ as required by the theory [20]. For our predictions of *π*_*T*_ under background selection, we calculated *G*_*ST, predicted*_ using background selection theory (see *Appendix*) and then computed *N*_*e, global*_. Using *N*_*e, global*_, we were then able to calculate the effect of background selection, *B*_*global*_, on *π*_*T*_ in the metapopulation. Then, we calculated predicted total diversity *π*_*T, pred*_ using the *tskit* package in Python 3.9.12 as:

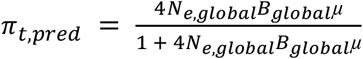

[43].

### Appendix: *F*_*ST*_ with a finite number of demes and migration immediately preceding measurement

Our simulations in SLiM involved a finite number of demes in an island model with drift preceding migration followed by assessment of the value of *F*_*ST*_. The effect of a finite number of demes has been assessed by several authors (see Ref. [32]); and having migration immediately precede assessment has been shown to reduce *F*_*ST*_ by Sved and Latter [37], albeit in a model with an infinitely large number of demes. Here we show calculations to predict *F*_*ST*_ of neutral loci accounting for both of these factors.

Here we will follow Slatkin [24], who showed that *F*_*ST*_ could be calculated as a function of the probability of nucleotide differences of two alleles chosen from the same sub-population (*π*_*S*_) or from two separate sub-populations (*π*_*B*_) in a finite island model with *d* demes. Slatkin [24] defined *F*_*ST*_ using the probability of sequence differences of two randomly chosen alleles, sampled without respect to which population they were from, or *π*_*T*_, which when all demes are equal in size is given by

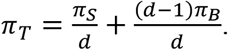

Slatkin’s definition of *F*_*ST*_ is

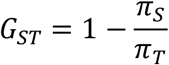

This definition of *F*_*ST*_ equivalent to what is often called *G*_*ST*_ (therefore we will refer to it as *G*_*ST*_ here to discriminate it from *F*_*ST*_ = *1* ™ π_*s*_/π_*B*_ that is estimated using π_*B*_ rather than π_*T*_ as the standard.)

Slatkin [24] showed that *F*_*ST*_ can be calculated from the mean coalescence times of a pair of alleles sampled both from the same population 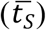 or from two separate populations 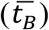.

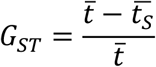

Here, t is the mean coalescent time of two alleles sampled at random without respect to which population they came from:

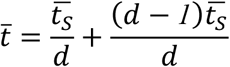

The equivalent expression for *F*_*ST*_ defined with π_*B*_ (as is implied, for example, in Weir and Cockerham’s estimate of *F*_*ST*_ [44]) is

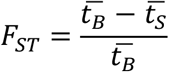

Slatkin showed that, for the case when migration precedes drift, the mean coalescent times for diploids are given by

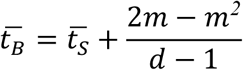

(Here we do not use the small migration approximation that Slatkin used for 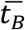) The latter equation calculates the time required for two alleles in different demes to have both been in the same deme in the last term, which then adds to the coalescent time of two alleles in the same deme for the final answer.

If the population is measured immediately after migration, then we can calculate the values of the mean coalescent times by modifying 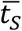 and 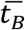 by one episode of migration. We will use asterisks to indicate the values of the mean coalescent times calculated in this way:

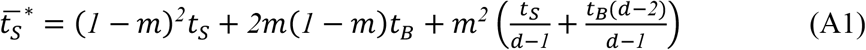

And

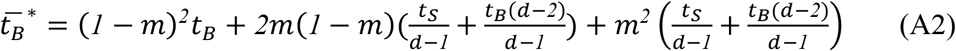

We can now find *F*_*ST*_ for this order of events:

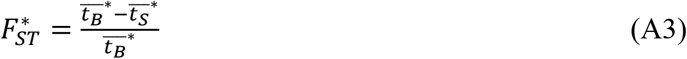

Putting equations A1 and A2 into Equation A3, we find, after some algebra:

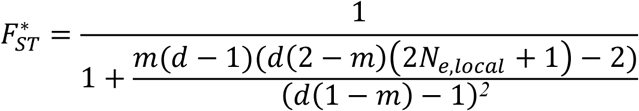

Assuming *N* >> 1, we can approximate this by

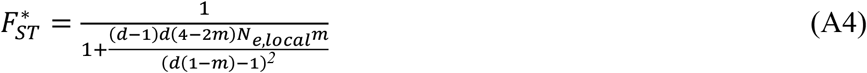

When the number of demes *d* is large, this recreates the result of Sved and Latter [37].

Let’s turn now to predicting *G*_*ST*_ immediately after migration, or

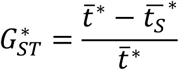

Using the values of t^*^ and <u>t_*s*_*</u> above we can find:

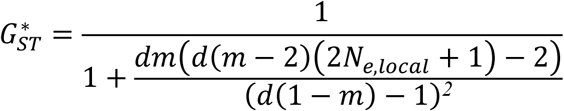

Again when *N* >> 1, this is well approximated by

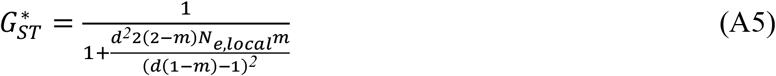

For haploids, the corresponding formula are

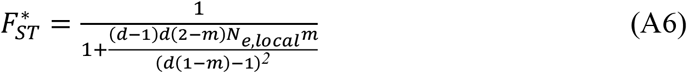

And

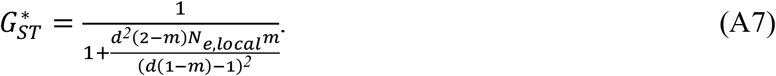

## Acknowledgements

We thank Tom Booker, Tyler Kent, Brian Charlesworth, and Tianlin Duan for helpful discussions and comments. Simulations for this study were conducted using the Zoology Computing Cluster supported by the Zoology Computing Unit at the University of British Columbia.

## Supporting Information

**Figure S1. Predicted *B* according to the Good et al. [29] method.** *B* was computed for haploid deleterious selection coefficients of *s* = {0.0005, 0.001, 0.002, 0.005, 0.01, 0.015, 0.03}, haploid population sizes of *N* = {100, 1000, 10000, 100000}, *U* = 7x10^-3^, and *R* = 0. Here we see that, as *N* increases, the theory predicts greater reductions in *B* for weaker selection coefficients (*s* ≤ 0.005).

**Figure S2. Predicted *π*_*S*_ versus observed *π*_*S*_ in a metapopulation.** (A) Forward-time simulations run with *m* = 0.00045 and *R* = 0, (B) *m* = 0.09 and *R* = 0, (C) *m* = 0.00045 and *R* = 0.01, and (D) *m* = 0.09 and *R* = 0.01. The gray dots represent forward-time simulations using a 10-deme WF Island model with local *N*_*e*_ = 1000, *N*_*local*_*s* = {0, 0.25, 0.5, 1, 2.5, 5, 10, 15}, *m* = {5x10^-5^, 1x10^-2^}, *r* = 0.005, and the purple dots represent theoretical predictions of total diversity (see *Methods*) with incorporation of the migration effect.

## Notes

### Competing Interest Statement

The authors have declared no competing interest.

